# Snail restricts SARS-CoV-2 entry by limiting ACE2 expression as part of broader remodelling of epithelial determinants of viral tropism

**DOI:** 10.64898/2026.10.20.760609

**Authors:** Prangya Paramita Sahoo, Vishal Sah, Krishan H Harshan

## Abstract

Pre-existing state of the host cell is a major determinant of the host susceptibility to viral infection. Here, we identify the epithelial-mesenchymal transition transcription factor Snail as a host restriction factor for SARS-CoV-2. Snail loss markedly increased SARS-CoV-2 replication, whereas Snail overexpression and complementation restricted viral replication. Transcriptomic analysis of Snail knockout cells revealed remodelling of epithelial surface-associated programs, including apical membrane, brush border, and junctional pathways. ACE2, the cellular receptor for SARS-CoV-2, was among the genes upregulated in Snail knockout cells. Snail negatively regulated *ACE2* transcription, leading to reduced ACE2 transcript and protein abundance and, consequently, lower SARS-CoV-2 entry. Consistent with this, *ACE2* promoter activity was higher in Snail knockout cells, which was reduced following Snail complementation. As a means to counter Snail’s antiviral functions, SARS-CoV-2 infection progressively depleted Snail protein in different cell culture models, while *SNAI1* transcript abundance was surprisingly maintained or increased under the same conditions. Proteasome inhibition and protein-turnover analyses failed to account for this depletion. We find that SARS-CoV-2 infection resulted in reduced association of *SNAI1* with polysomes at later stages of infection coinciding with virus-induced host translational suppression, resulting in lower protein synthesis. Together, these findings identify Snail as a regulator of the epithelial entry landscape that limits ACE2-associated SARS-CoV-2 entry and shapes cellular permissiveness, with broader implications for cellular determinants of viral tropism. SARS-CoV-2, in turn, progressively depletes this short-lived antiviral regulator, revealing how virus-induced translational shutoff can remodel pre-existing host cell states and thereby alter cellular susceptibility during infection.

**IMPORTANCE:** Cellular permissiveness is a major determinant of viral tropism. Host transcriptional programs are well known to shape cellular identity and surface composition, but their contribution to viral susceptibility is less often viewed as a form of host restriction. We previously showed that the epithelial-mesenchymal transition transcription factors Snail and Slug restrict multiple RNA viruses. Here, we show that Snail also limits SARS-CoV-2 infection by restricting ACE2 expression and by remodelling the epithelial surface landscape. SARS-CoV-2 neutralizes this threat by translationally repressing *SNAI1* as part of the global translation suppression. We show that loss of Snail remodels surface-associated cellular programs and establishes a more permissive state. Together, these findings connect two aspects of host-virus interaction: transcriptional programs that establish cellular permissiveness and viral processes that can subsequently alter them. They position Snail as a broader regulator of the cellular landscape that influences viral entry and tropism.

## INTRODUCTION

Viruses rely extensively on host-cell factors to establish productive infection. Severe acute respiratory syndrome coronavirus 2 (SARS-CoV-2) entry is initiated by the engagement of Spike glycoprotein with angiotensin-converting enzyme 2 (ACE2), followed by proteolytic activation of Spike by host protease such as TMPRSS2 (1). Although other cell surface receptors are identified for facilitating SARS-CoV-2 entry, ACE2 abundance and its accessibility at the cell surface represent important determinants of cellular susceptibility to SARS-CoV-2 (1–3). Viral tropism at the cellular level is primarily determined by the presence or absence of entry receptors and other host factors (4–6). SARS-CoV-2 displays favourable tropism towards epithelial cells (7, 8), suggesting that changes in epithelial cell state could alter the host-factor landscape encountered by the virus and thereby influence cellular permissiveness and tropism.

Epithelial-to-mesenchymal transition (EMT) is a phenomenon in which epithelial cells lose their characteristics and acquire those of mesenchymal cells driven by major transcriptional reprogramming (9, 10). This transition involves coordinated changes in polarity, cell-cell adhesion, cytoskeletal organization, and expression of epithelial surface proteins (11). Several epithelial markers and junctional proteins, including E-cadherin, Claudins, and Occludin, are reduced during EMT (9), driven by transcription factors belonging primarily to the SNAIL, ZEB, and TWIST families.

Changes induced by these factors can have uniquely different consequences for individual viruses depending on their host-factor requirements. Snail induced EMT reduces susceptibility of polarized epithelial cells to measles virus by altering an epithelial receptor associated program, whereas EMT can enhance herpes simplex virus type 1 infection through increased expression of its entry receptor nectin-1 (12, 13). Thus, EMT can either restrict or facilitate infection depending on how the resulting cellular state alters receptors and other virus-supporting host factors.

Interestingly, studies indicate that ACE2 expression is also influenced during EMT. In lung epithelial and cancer models, induction of EMT, including activation of the EMT transcription factor ZEB1, is associated with repression of *ACE2*, establishing a link between epithelial state and SARS-CoV-2 susceptibility (14). Such changes in epithelial identity can alter cellular permissiveness and, consequently, viral tropism by modifying the availability of receptors and other host factors required for infection.

Snail, encoded by *SNAI1*, is a zinc-finger transcription factor that recognizes E-box-containing regulatory sequences and is best characterized for repressing epithelial genes during EMT (15). In SARS-CoV-2 models, Snail induction and EMT-like changes have also been reported, although the functional consequence of Snail during productive SARS-CoV-2 infection remains unclear (16). Although Snail has traditionally been studied in the context of development, fibrosis, and cancer, emerging evidence indicates that EMT transcription factors can also participate in cell-intrinsic antiviral responses. Our previous work demonstrated that RNA virus infection induces *SNAI1* transcription as part of the host antiviral response.

Increased Snail expression, in turn, was associated with enhanced antiviral signaling and restricted the replication of multiple RNA viruses, including dengue virus, Japanese encephalitis virus, and vesicular stomatitis virus, whereas Snail depletion enhanced viral replication (17). These findings identified Snail as an antiviral regulator and suggested that EMT-associated transcription factors can have functions during infection that are distinct from their conventional roles in cellular plasticity. However, whether Snail similarly restricts SARS-CoV-2, and whether such activity is primarily linked to its previously described relationship with interferon signaling or to an alternative antiviral mechanism, has not been established.

In this study, we investigated the antiviral function and regulation of Snail during SARS-CoV-2 infection. We find that Snail exhibits potent antiviral functions against SARS-CoV-2 where the loss of Snail resulted in remarkably elevated levels of viral replication. Interestingly, this antiviral activity did not involve IFN signalling, but was imparted through reducing the viral permissiveness of the cells by remodelling epithelial cell surface gene programs. Notably, ACE2, the major entry receptor of SARS-CoV-2, was among the transcriptionally suppressed genes by Snail. Our results indicate that Snail uses this mechanism to regulate viral entry, thereby reducing viral permissiveness. Whereas Snail exhibits strong antiviral effects, we show that SARS-CoV-2 targets Snail expression through translational suppression, thereby neutralizing the antiviral functions of Snail. Our study demonstrates how fundamental characteristics of the host cell dictate viral permissiveness and EMT-TFs can alter viral tropism by changing these cellular features.

## RESULTS

### Snail limits SARS-CoV-2 infection

We previously reported that the EMT transcription factors Snail and Slug exert antiviral activity against multiple RNA viruses (17). SARS-CoV-2 unusually delays the activation of IFN response unlike many other viruses (18). We therefore asked if Snail’s antiviral effects are extended to limit SARS-CoV-2 infection. Despite repeated attempts, we failed to observe appreciable knockdown (KD) of Snail or to generate Snail KO in Caco-2 cells using CRISPR-Cas9 approach. Nevertheless, we successfully generated Snail KO Huh7 cells. These epithelial like cells of hepatic origin express ACE2 and support robust productive replication of SARS-CoV-2 (19–21). We measured SARS-CoV-2 infection in both pooled and clonal cultures of Snail KO Huh7 cells. Complete absence of Snail was evident in the two independently derived KO pool cultures while it was robustly detected in the WT cells (Supplementary Figure S1A). Snail KD resulted in significantly elevated SARS-CoV-2 replication, as evident from higher N levels and viral titers (∼ 2-3 logs increase) in two independent KD pool cultures (Supplementary Figure S1 A-C). We further confirmed this observation in clonally selected Snail KO cells. Snail loss was verified by immunoblotting (Figure 1A). Similar to the KD data, substantially higher amounts of Viral Nucleocapsid (N) protein were detected in the Snail KO cells from a more robust viral replication (∼ 2 logs) as compared with the WT cells (Figure 1 A-C), validating the antiviral effects of Snail against SARS-CoV-2. We further confirmed this finding in the converse experiment, where we overexpressed Snail in Huh7 cells followed by SARS-CoV-2 infection. Robust Snail overexpression resulted in marked drop in N levels and ∼ 2 log drop in viral titers (Figures 1 D-F). To further verify that the enhanced infection in Snail KO cells was attributable specifically to the loss of Snail rather than to an off-target consequence of clonal selection, we re-expressed Snail in Snail KO cells. Complementation of Snail in the KO cells markedly reduced N levels and the viral titers (Figures 1 G-I), underlining the antiviral functions of Snail against SARS-CoV-2.

**Figure 1.**
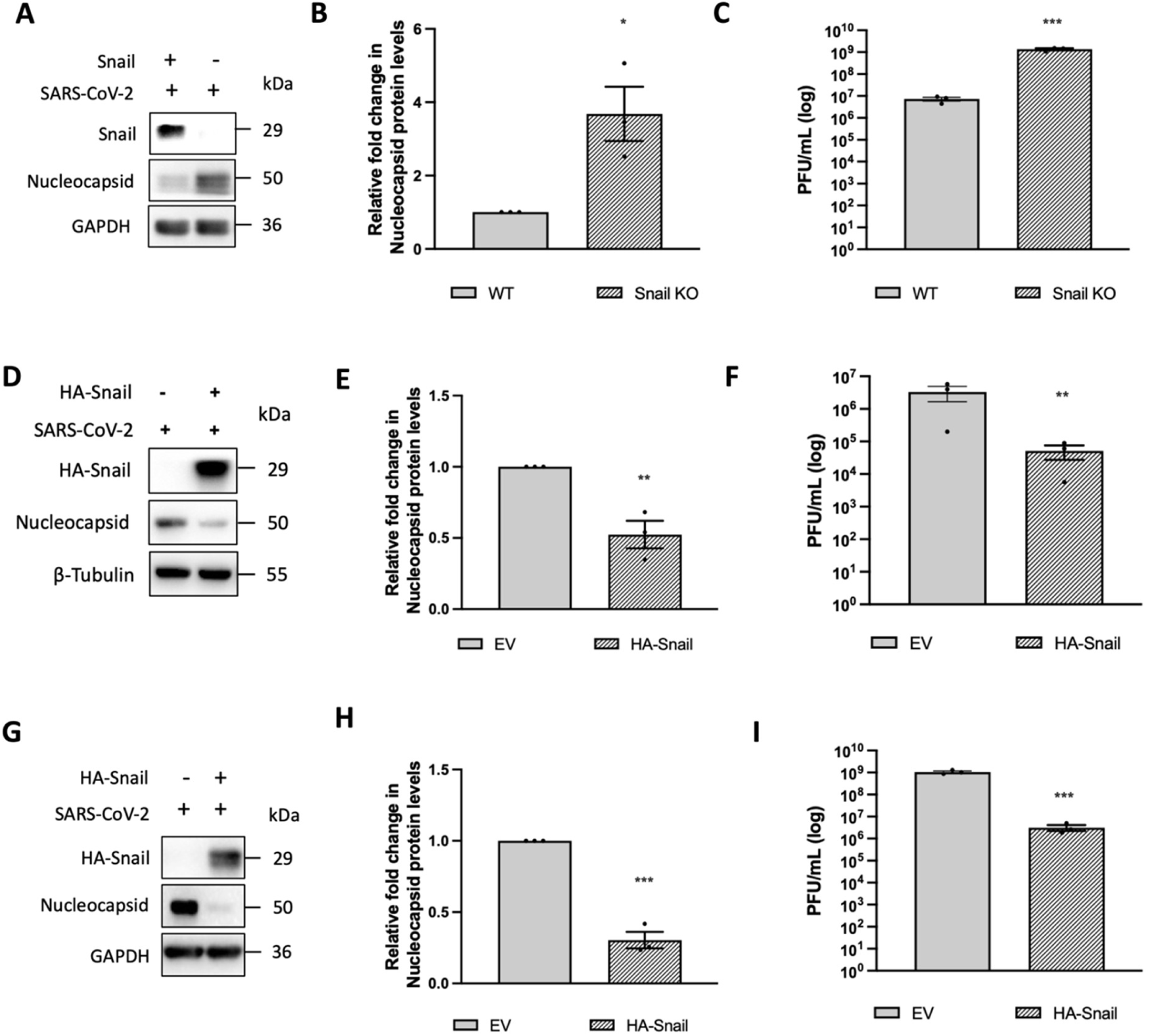
Snail limits SARS-CoV-2 infection. (A) Immunoblot showing Snail, SARS-CoV-2 Nucleocapsid, and GAPDH protein levels in WT and Snail KO Huh7 cells following SARS-CoV-2 infection. (B) Densitometric quantification of Nucleocapsid protein levels from panel (A). Individual Nucleocapsid band intensities from both the WT and Snail KO samples were normalized to their corresponding GAPDH band intensities. The normalized values from the Snail KO cells were plotted relative to those from the WT cells. (C) Infectious viral titre in SARS-CoV-2 infected WT and Snail KO cells, represented as PFU/mL. (D) Immunoblot showing HA-Snail, SARS-CoV-2 Nucleocapsid, and β-Tubulin protein levels in SARS-CoV-2-infected Huh7 cells transfected with pcDNA4/TO empty vector or HA-Snail-pcDNA4/TO. (E) Densitometric quantification of Nucleocapsid protein levels from panel (D). Individual Nucleocapsid band intensities from the pcDNA4/TO empty vector or HA-Snail-pcDNA4/TO transfected samples were normalized to their corresponding β-Tubulin band intensities. The values for HA-Snail-pcDNA4/TO were plotted relative to those from the pcDNA4/TO transfected samples. (F) Infectious viral titre in SARS-CoV-2-infected Huh7 cells transfected with pcDNA4/TO or HA-Snail-pcDNA4/TO, represented as PFU/mL. (G) Immunoblot showing HA-Snail, SARS-CoV-2 Nucleocapsid, and GAPDH protein levels in Snail KO cells transfected with pcDNA4/TO or HA-Snail-pcDNA4/TO following SARS-CoV-2 infection. (H) Densitometric quantification of Nucleocapsid protein levels from panel (G). Individual Nucleocapsid band intensities from the pcDNA4/TO empty vector or HA-Snail-pcDNA4/TO transfected samples were normalized to their corresponding GAPDH band intensities. The values for HA-Snail-pcDNA4/TO were plotted relative to those from the pcDNA4/TO transfected samples (I) Infectious viral titre in SARS-CoV-2 infected Snail KO cells transfected with pcDNA4/TO empty vector or HA-Snail-pcDNA4/TO represented as PFU/mL. Data are represented as mean ± SEM from three independent biological replicates (n = 3). Each dot represents one biological replicate. Statistical significance was determined using an unpaired Student’s *t*-test. *, **, and *** denote *P* < 0.05, *P* < 0.01, and *P* < 0.001, respectively; n.s., not significant.

IFN signaling displayed limited restriction of SARS-CoV-2 variants emerged later as against the earlier variants against which it showed robust restriction (22). Given these differential responses against IFN, we tested if Snail-mediated antiviral effects generated broad antiviral state. Since we used B.1.1.8 isolate, a variant isolated early during COVID-19 pandemic, in this study so far, we tested its efficacy against a Delta variant isolate. As shown in Supplementary Figure S1 D-F, Snail’s antiviral potency was retained against this later variant, demonstrating that Snail elicits a broad antiviral response and possibly uses mechanisms that are shared by the distinct SARS-CoV-2 variants.

### SARS-CoV-2 infection progressively depletes Snail protein despite maintained *SNAI1* transcript abundance

Viral infections frequently remodel or antagonize host restriction factors to create a more permissive intracellular environment (23, 24). Having established that Snail restricts SARS-CoV-2 infection, we next asked whether the virus alters the abundance of this antiviral host factor during infection as reported in our previous article (17). To determine how Snail expression is regulated during SARS-CoV-2 infection, we first examined its abundance in Huh7 cells over the course of infection. Huh7 cells are hepatoma cell lines with epithelial characteristics and show high SARS-CoV-2 permissiveness (21, 25). Cells were infected with 1 MOI of SARS-CoV-2 and harvested at 12-, 24-, 48-, and 72 hpi (Figure 2A). N protein was readily detected in the infected cells and progressively accumulated with infection, confirming productive infection (Figure 2B). Snail protein abundance remained largely stable until 12 hpi, but surprisingly underwent sharp decline starting from 24 hpi (Figures 2 B&C). A pronounced reduction in Snail level was evident from 48 hpi onwards.

**Figure 2.**
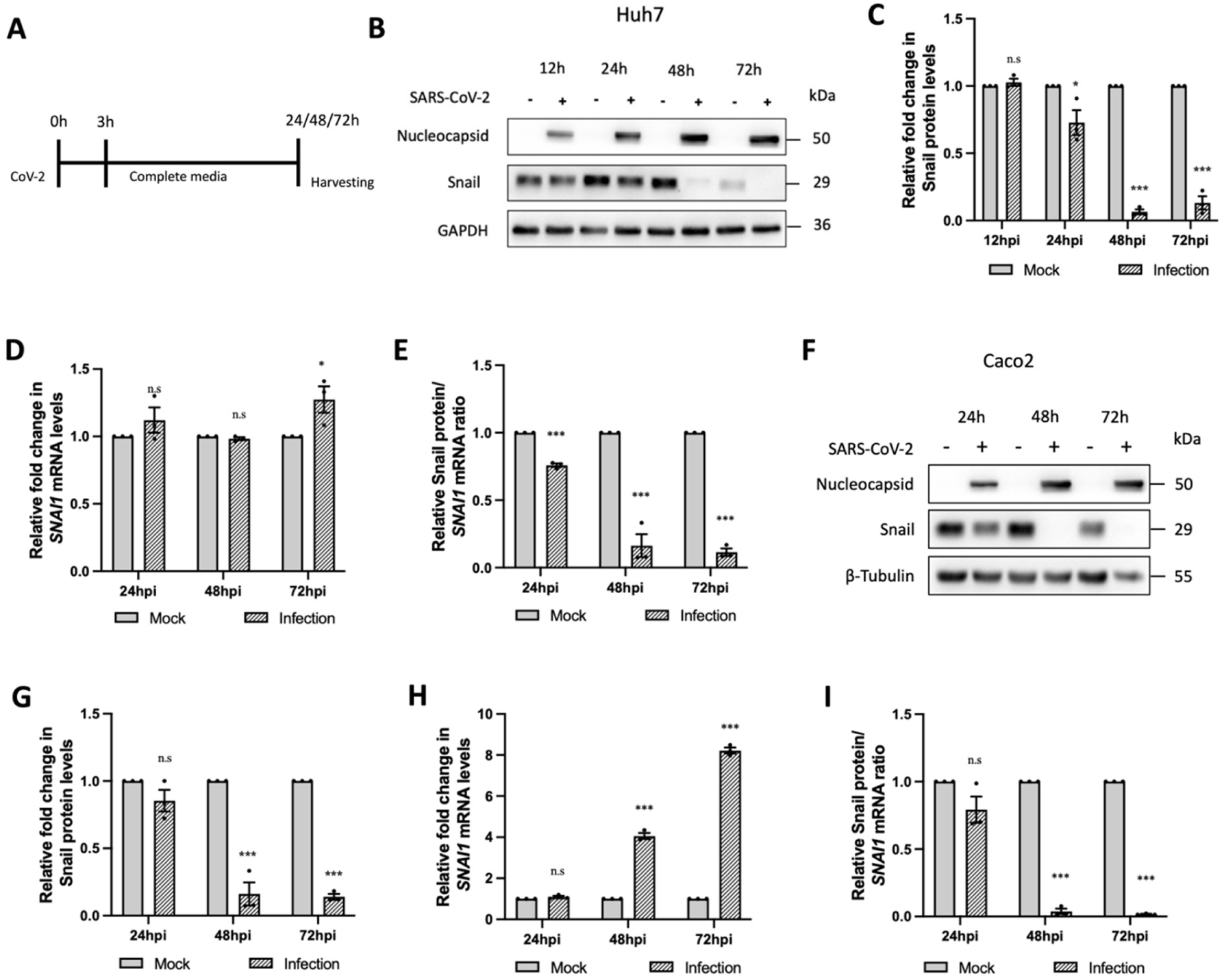
SARS-CoV-2 infection causes progressive Snail protein loss despite maintained *SNAI1* transcript abundance. (A) Schematic representation of the SARS-CoV-2 infection timeline in Huh7 cells. Cells were infected for 3 hrs, followed by replacement of the inoculum with complete medium and harvesting at the indicated time points. (B) Immunoblot showing SARS-CoV-2 Nucleocapsid, Snail, and GAPDH protein levels in mock and SARS-CoV-2-infected Huh7 cells at 12-, 24-, 48-, and 72 hpi. (C) Densitometric quantification of Snail protein levels from panel (B). Individual Snail band intensities from the mock and SARS-CoV-2-infected samples were normalized to their corresponding GAPDH band intensities. The normalized values from the infected cells were plotted relative to those from the corresponding time-matched mock controls. (D) RT-qPCR analysis of *SNAI1* mRNA expression in mock and SARS-CoV-2-infected Huh7 cells at 24-, 48-, and 72 hpi. (E) Plot representing relative Snail protein:*SNAI1* mRNA ratio during SARS-CoV-2 infection in Huh7 cells. The ratio was calculated by dividing the normalized Snail protein abundance by the normalized *SNAI1* transcript abundance at each time point. (F) Immunoblot showing SARS-CoV-2 Nucleocapsid, Snail, and β-Tubulin protein levels in mock and SARS-CoV-2-infected Caco-2 cells at 24-, 48-, and 72 hpi. (G) Densitometric quantification of Snail protein levels from panel (F). Individual Snail band intensities from the mock and SARS-CoV-2-infected samples were normalized to their corresponding β-Tubulin band intensities. The normalized values from the infected cells were plotted relative to those from the corresponding time-matched mock controls. (H) RT-qPCR analysis of *SNAI1* mRNA expression in mock and SARS-CoV-2-infected Caco-2 cells at 24-, 48-, and 72 hpi. (I) Plot representing relative Snail protein:*SNAI1* mRNA ratio during SARS-CoV-2 infection in Caco-2 cells. The ratio was calculated by dividing the normalized Snail protein abundance by the normalized *SNAI1* transcript abundance at each time point. Data are represented as mean ± SEM from three independent biological replicates (n = 3). Each dot represents one biological replicate. Statistical significance was determined using an unpaired Student’s *t*-test. *, **, and *** denote *P* < 0.05, *P* < 0.01, and *P* < 0.001, respectively; n.s., not significant.

Since *SNAI1* transcription is subject to upstream regulation including during viral infections (16, 17), we measured *SNAI1* levels in these samples by using RT-qPCR. In contrast to the progressively dropping Snail levels, *SNAI1* transcript levels were maintained at 24- and 48-hpi and then spiked at 72 hpi in Huh7 cells (Figure 2D).

Consequently, the relative Snail protein:*SNAI1* mRNA ratio progressively declined over the course of infection (Figure 2E), indicating increasing uncoupling between *SNAI1* transcript abundance and Snail protein levels. Since RT-qPCR does not necessarily discriminate between full-length and shorter products of transcripts, we compared the full-length *SNAI1* transcripts in the infected Huh7 cells by northern blotting. As shown in Supplementary Figures 2 A&B, *SNAI1* transcript levels were preserved throughout the course of infection, confirming that SARS-CoV-2 does not target *SNAI1* transcription.

To determine whether this phenomenon is cell type-specific, we examined Snail regulation in additional epithelial cell line models. Strong depletion of Snail was consistently observed in Caco-2, a colon epithelial cell line highly permissive to SARS-CoV-2 (Figures 2 F&G), and A549, a lung adenocarcinoma cell line with relatively lower SARS-CoV-2 permissiveness; (Supplementary Figures 2 C&D). N protein abundance remained strong throughout the infection, indicating that the depletion is specifically targeting Snail. These results suggested that SARS-CoV-2 infection progressively depletes Snail protein across epithelial cell models. Notably, SARS-CoV-2 infection in Caco-2 and A549 also resulted in marked increase in *SNAI1* transcript levels early in infection that progressively increased with the infection (Figure 2H and Supplementary Figure 2E). The marked decrease in the relative Snail protein:*SNAI1* mRNA ratio over the course of infection in these two additional cell models (Figure 2I and Supplementary Figure 2F) confirmed that SARS-CoV-2 targets Snail levels through a post-transcriptional mechanism without involving the stability of *SNAI1* transcript.

### SARS-CoV-2-induced host translational shutdown drives the depletion of short-lived Snail protein

Since the Snail depletion coincides with the delayed activation of type I IFNs in SARS-CoV-2 infected cells (22, 26), we asked if Snail depletion is influenced by IFN signaling. To investigate this, we pre-treated Huh7 cells with IFNα prior to SARS-CoV-2 infection and maintained the infected cells in IFNα containing medium until harvested at 24 hpi, a time point in infection before IFN signaling is activated (Figure 3A). IFNα treatment induced robust STAT1 phosphorylation and IFIT1 expression as expected, confirming the activation of IFN signaling (Figure 3B). Consistent with the establishment of antiviral state, N levels were reduced in IFNα treated samples.

**Figure 3.**
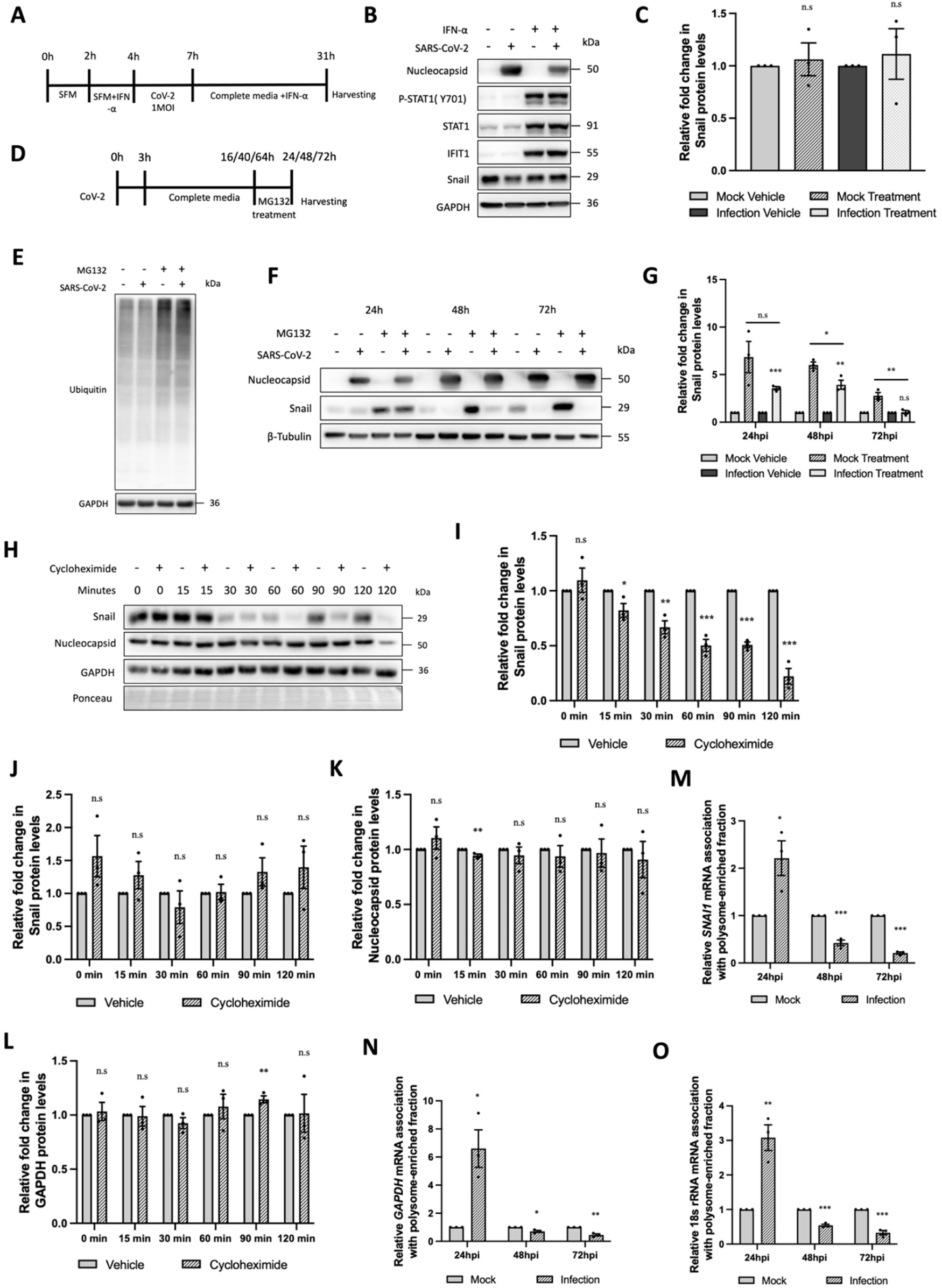
SARS-CoV-2-induced host translational shutdown drives the depletion of short-lived Snail protein. (A) Schematic representation of the IFN-α treatment experiment. Huh7 cells were maintained in serum-free DMEM for 3 hrs, followed by pretreatment with 1,000 U/mL IFN-α for 1 hr before SARS-CoV-2 infection. Following 3 hrs of infection, the inoculum was replaced with complete medium containing IFN-α, and cells were maintained until harvest at 24 hpi. (B) Immunoblot showing SARS-CoV-2 Nucleocapsid, p-STAT1 (Y701), total STAT1, IFIT1, Snail, and GAPDH protein levels in mock and SARS-CoV-2-infected Huh7 cells treated with vehicle or IFN-α. (C) Densitometric quantification of Snail protein levels from panel (B). Individual Snail band intensities from mock-and SARS-CoV-2-infected cells treated with vehicle or IFN-α were normalized to their corresponding GAPDH band intensities. The normalized values were plotted relative to the mock vehicle-treated control. (D) Schematic representation of the MG132 treatment experiment. Huh7 cells were infected with SARS-CoV-2 for 3 hrs, followed by replacement with complete medium. MG132 was added 8 h before harvesting at 24-, 48-, and 72 hpi. (E) Immunoblot showing ubiquitinated proteins and GAPDH in mock-and SARS-CoV-2-infected Huh7 cells treated with vehicle or MG132. (F) Immunoblot showing SARS-CoV-2 Nucleocapsid, Snail, and β-Tubulin protein levels in mock- and SARS-CoV-2-infected Huh7 cells treated with vehicle or MG132 at 24-, 48-, and 72 hpi. (G) Densitometric quantification of Snail protein levels from panel (F). Individual Snail band intensities from mock-and SARS-CoV-2-infected cells treated with vehicle or MG132 were normalized to their corresponding β-Tubulin band intensities. At each time point, the normalized values for all conditions were plotted relative to the corresponding mock vehicle-treated control. (H) Cycloheximide chase analysis of Snail protein stability in SARS-CoV-2-infected Huh7 cells. Snail, GAPDH, and SARS-CoV-2 Nucleocapsid protein levels were analyzed at the indicated times following cycloheximide or vehicle treatment. Ponceau staining was used to assess total protein loading. (I) Densitometric quantification of Snail protein levels from panel (H). Individual Snail band intensities were normalized to their corresponding Ponceau total-protein signals. At each time point, the normalized value from the cycloheximide-treated sample was plotted relative to its corresponding time-matched vehicle control. (J) Plot showing the comparison of Snail protein turnover between mock- and SARS-CoV-2-infected cells following cycloheximide treatment. Snail remaining after cycloheximide treatment was first calculated relative to the corresponding time-matched vehicle control within each condition. The normalized value from the infected cells was then divided by the corresponding mock value at each time point and represented as the infected/mock ratio. (K) Densitometric quantification of SARS-CoV-2 Nucleocapsid protein levels from panel (H). Individual Nucleocapsid band intensities were normalized to their corresponding Ponceau total-protein signals. At each time point, the normalized value from the cycloheximide-treated sample was plotted relative to its corresponding time-matched vehicle control. (L) Densitometric quantification of GAPDH protein levels from panel (H). Individual GAPDH band intensities were normalized to their corresponding Ponceau total-protein signals. At each time point, the normalized value from the cycloheximide-treated sample was plotted relative to its corresponding time-matched vehicle control. (M-O) Polysome-enriched pellet and non-polysomal supernatant fractions were collected from mock and SARS-CoV-2-infected Huh7 cells at 24-, 48-, and 72 hpi, followed by RNA isolation and RT-qPCR analysis. Relative pellet to supernatant ratios are shown for (M) *SNAI1* mRNA, (N) *GAPDH* mRNA, and (O) 18S rRNA. The ratios from SARS-CoV-2-infected samples were plotted relative to the corresponding time-matched mock controls. Data are represented as mean ± SEM from three independent biological replicates (n = 3). Each dot represents one biological replicate. Statistical significance was determined using an unpaired Student’s *t*-test. *, **, and *** denote *P* < 0.05, *P* < 0.01, and *P* < 0.001, respectively; n.s., not significant.

However, Snail protein abundance remained unaffected by IFNα treatment in both SARS-CoV-2 infected and uninfected samples, suggesting that IFN signaling does not influence Snail abundance in our experimental set up (Figure 3 B&C).

Next, we considered different mechanisms of post-transcriptional Snail regulation during SARS-CoV-2 infection. Snail abundance is known to be regulated through several post-transcriptional mechanisms including rapid protein turnover through the ubiquitin-proteasome system (UPS) (27). Further, SARS-CoV-2 is well reported to hijack host UPS to selectively target host proteins (28). We first addressed if Snail degradation during the infection is mediated through proteasomal degradation. Huh7 cells infected with SARS-CoV-2 were treated with proteasome inhibitor MG132 for 8 hrs before harvesting at 24-, 48-, or 72 hpi (Figure 3D). Accumulation of ubiquitinated proteins following the treatment confirmed effective inhibition of proteasomal activity (Figure 3E). As expected for a protein subject to rapid turnover, proteasome inhibition resulted in increased basal Snail abundance in mock-infected cells (Figures 3 F&G). However, MG132 failed to restore Snail protein abundance in SARS-CoV-2 infected cells, particularly in the later stages of infection when Snail depletion was prominent. Thus, although proteasomal turnover contributes to steady-state Snail abundance, enhanced proteasomal degradation alone looks insufficient to explain the profound Snail depletion seen in SARS-CoV-2 infection. SARS-CoV-2 main protease (Nsp5) is known to target several host proteins for proteolytic cleavage (29, 30). We next asked if Snail is targeted by viral proteases. In silico sequence analysis identified a putative Nsp5 cleavage motif in Snail sequence at aa residue 149 generating two polypeptide products of 16 and 13 kDa respectively (Supplementary Figure S3A). However, immunoblot analysis failed to detect the two cleaved polypeptide products of Snail in SARS-CoV-2 infected samples (Supplementary Figure S3B), ruling out the possibility of Nsp5 mediated cleavage of Snail behind its depletion during the infection.

As Snail is a short-lived protein, we asked whether SARS-CoV-2 infection expedites its turnover rate. Cycloheximide (CHX) chase assay in mock infected Huh7 cells showed rapid turnover of Snail starting from 30 minutes of CHX treatment (Supplementary Figures 3 C&D). We then performed the same assay in SARS-CoV-2-infected cells. A similar progressive decline in Snail was observed in SARS-CoV-2-infected cells following CHX treatment (Figures 3 H&I). We therefore directly compared Snail turnover between the mock and the infected conditions to determine whether infection accelerated this process. For this comparison, normalized Snail levels in infected cells were plotted relative to the corresponding mock values at each time point (infected/mock ratio; Figure 3J). If SARS-CoV-2 infection accelerated Snail degradation, this ratio is expected to decrease during the CHX chase.

However, no consistent decline in this ratio was observed over the course of the experiment, indicating that SARS-CoV-2 infection did not substantially accelerate Snail turnover. By contrast, SARS-CoV-2 Nucleocapsid and GAPDH protein abundance remained largely unchanged during the CHX chase, confirming the specificity of this observation (Figures 3 K&L). Similar observations were made in GAPDH abundance in the mock-infected samples as well (Supplementary Figure 3E).

Our results so far demonstrate that SARS-CoV-2-mediated Snail depletion is a post-transcriptional event, independent of proteasomal degradation and proteolytic cleavage. This prompted us to consider the possibility of translational regulation of *SNAI1* transcript during SARS-CoV-2 infection. SARS-CoV-2 is reported to force a strong global translation suppression primarily through Nsp1 (31, 32). We studied the translational engagement of *SNAI1* transcript in SARS-CoV-2 infected cells by comparing its polysomal association. Polysomes were precipitated from both the SARS-CoV-2 infected and uninfected samples collected at 24-, 48-, and 72 hpi, and *SNAI1* transcripts were quantified in the polysome enriched and non-polysome supernatant fractions. The ratio of *SNAI1* transcript abundance between the polysome-enriched pellet and non-polysomal supernatant fractions was plotted to assess its polysome association. Figure 3M demonstrated a higher *SNAI1* mRNA:polysome association at 24 hpi, before showing a sharp drop at 48 hpi onwards. This result indicated an active and continuous synthesis of Snail protein until 24 hpi, but a sharp decline in its synthesis at later time points, as reflected in the immunoblots (Figure 2B). To determine whether this reduced polysome association was specific to *SNAI1* mRNA or reflected a broader disruption of host translation, we also examined *GAPDH* mRNA, which is unrelated to Snail and a housekeeping transcript. *GAPDH* mRNA showed a similar reduction in polysome association at later stages of infection, arguing against a *SNAI1*-specific translation suppression (Figure 3N). We also quantified 18S rRNA as a ribosome marker to assess whether the changes in mRNA association were accompanied by changes in ribosome association with the polysome-enriched fraction. 18S rRNA also showed a decline at 48 and 72 hpi (Figure 3O), further supporting a broader disruption of host translation during infection. Together, these findings suggest that reduced translational engagement of *SNAI1* transcript during SARS-CoV-2 infection contributes to the progressive depletion of the short-lived Snail protein despite maintained *SNAI1* transcript abundance.

### Snail depletion remodels the epithelial entry landscape and enhances SARS-CoV-2 entry

The antiviral effect of Snail prompted us to investigate the cellular mechanism underlying this restriction and the consequences of Snail depletion on SARS-CoV-2 infection. Since Snail has previously been associated with antiviral innate immune signaling and IFNs are the key antiviral players in innate antiviral response, we first examined whether the enhanced infection observed following Snail depletion could be explained by altered IFN production or responsiveness. Overexpression of Snail in Huh7 cells did not significantly alter the transcript abundance of *IFNB1* and *IFIT1,* important indicators of type I IFN production and downstream IFN signaling (Supplementary Figures S4 A&B). No change in their levels was also detected in SARS-CoV-2 infection, confirming their non-involvement in Snail’s antiviral functions. *IFNL1* transcripts also remained unaffected under these conditions, ruling out the involvement of type III IFN signaling in the process (Supplementary Figure S4C). We next sought to test if the IFN responsive pathways are intact in Snail KO cells during SARS-CoV-2 infection. IFNα treatment activated STAT1 phosphorylation in both Snail KO and their WT counterpart cultures infected with SARS-CoV-2 at equal efficiency, (Supplementary Figure S4D). Additionally, comparable drop in N levels and infectious viral titres was observed in IFNα treated Snail KO and WT cells, further confirming that IFN is not a mediator in Snail’s antiviral functions (Supplementary Figures S4 D-F). We therefore investigated alternate cellular pathways that could account for Snail’s antiviral effects.

To obtain an unbiased view of cellular pathways altered following Snail depletion, we compared the transcriptomic profiles of WT and Snail KO cells under both mock and SARS-CoV-2 infected conditions. Previous studies using Snail KO models have established a broad transcriptional reprogramming using differential gene expression analysis (33). The differential expression analysis between WT and Snail KO Huh7 cells revealed broad transcriptional changes in Snail KO cells under both the infected and uninfected conditions (Supplementary Figure S4G). We next compared the differentially expressed genes (DEGs) identified in Snail KO versus WT cells under mock and SARS-CoV-2-infected conditions using a Venn analysis (Supplementary Figure S4H). Of the 604 DEGs identified across the two comparisons, 297 genes (49.2%) were shared between mock and infected conditions, whereas 129 genes were unique to the mock condition and 178 were unique to the infected condition.

The large shared fraction suggests that a substantial part of the transcriptional changes attributed to Snail loss remained unaffected during SARS-CoV-2 infection. We therefore focused on these commonly altered genes to identify Snail dependent cellular pathways that could define the pre-existing cellular state encountered by SARS-CoV-2. Gene ontology (GO) analysis of these commonly altered genes revealed enrichment of terms associated with epithelial organization and membrane architecture including apical plasma membrane, brush border, apical junction complex, and epithelial surface associated processes (Supplementary Figure S4I).

Since Snail depletion increased SARS-CoV-2 permissiveness, we next focused on genes that were upregulated in Snail KO cells, reasoning that increased expression of host factors or cellular programs that favor viral susceptibility could contribute to this phenotype. Consistent with the enrichment observed among the commonly altered genes, GO analysis of the upregulated gene set again highlighted epithelial surface associated pathways, including apical plasma membrane, brush border, and cell-cell junction organization (Figure 4A). The enrichment of epithelial surface associated pathways suggested that Snail loss remodels the cellular interface encountered by the virus, potentially influencing viral tropism through changes in the availability and organization of entry associated host factors (34–36). We therefore examined individual genes contributing to these enriched pathways. *ACE2* was identified among the upregulated genes associated with the apical plasma membrane category (Figure 4B) and was also represented within the brush border and cell-cell junction organization gene sets (Supplementary Figures S4 J&K). The recurrent identification of *ACE2* across these independent enrichment categories was particularly interesting given its established role as the primary cellular receptor for SARS-CoV-2 entry (1, 37).

**Figure 4.**
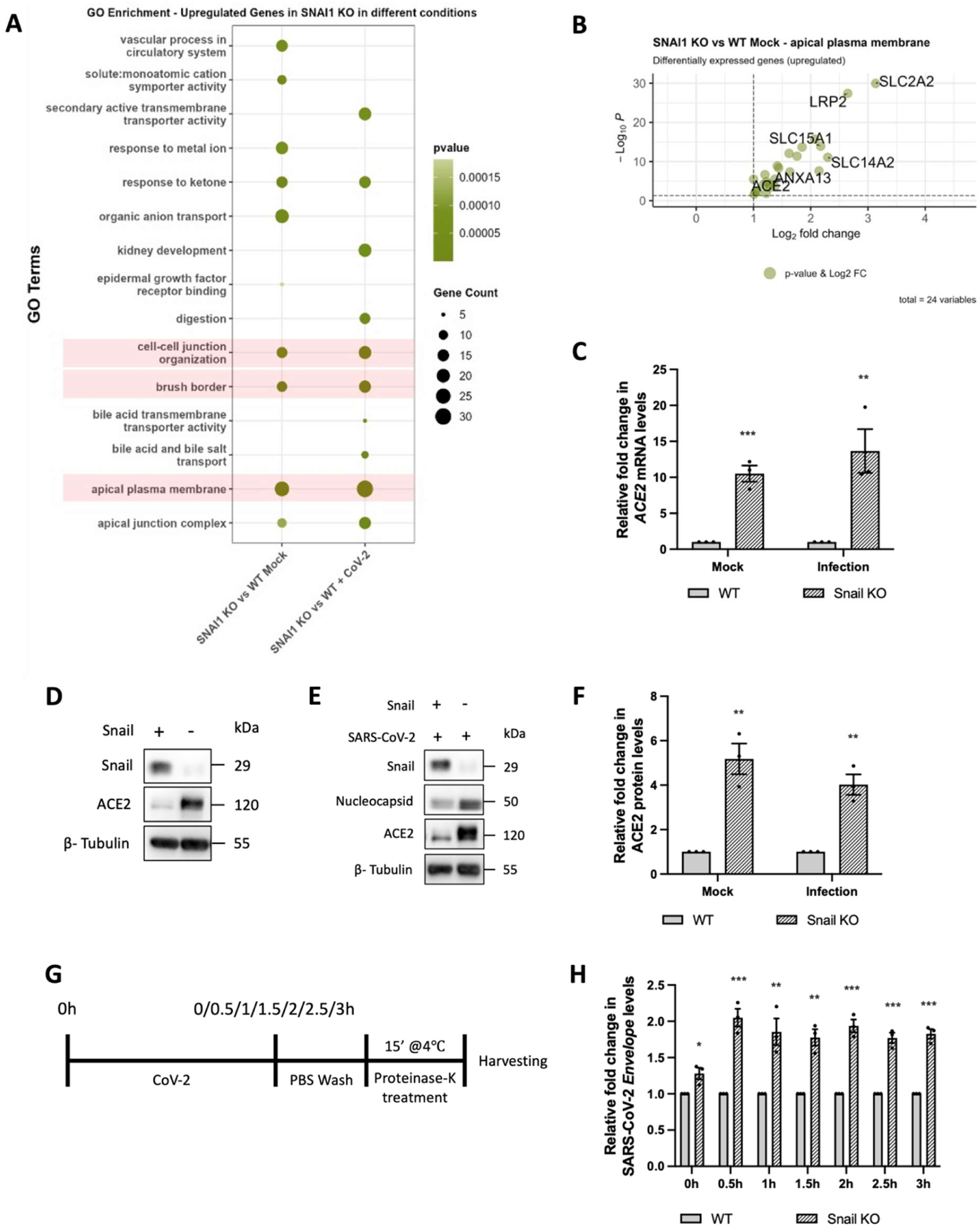
Snail depletion remodels the epithelial entry landscape and enhances SARS-CoV-2 entry. (A) Gene ontology enrichment analysis of genes upregulated in Snail KO cells compared with WT cells under mock and SARS-CoV-2-infected conditions. RNA deep seq was performed in WT and Snail KO Huh7 cells under both conditions. Gene counts were normalized using DESeq2, and differential gene expression was determined by comparing Snail KO with WT cells separately under mock and SARS-CoV-2-infected conditions. Genes upregulated in Snail KO cells from each comparison were subjected to GO enrichment analysis. Dot size represents gene count, and colour indicates adjusted *P* value. (B) Volcano plot showing upregulated genes associated with the apical plasma membrane gene ontology term in uninfected Snail KO cells compared with uninfected WT cells. (C) RT-qPCR analysis of *ACE2* mRNA expression in WT and Snail KO cells under mock- and SARS-CoV-2-infected conditions. The normalized values from the Snail KO cells were plotted relative to those from the corresponding WT control under each condition. (D) Immunoblot showing Snail, ACE2, and β-Tubulin protein levels in uninfected WT and Snail KO Huh7 cells. (E) Immunoblot showing Snail, SARS-CoV-2 Nucleocapsid, ACE2, and β-Tubulin protein levels in WT and Snail KO cells following SARS-CoV-2 infection. (F) Densitometric quantification of ACE2 protein levels from panels (D) and (E). Individual ACE2 band intensities from WT and Snail KO samples under mock- and SARS-CoV-2-infected conditions were normalized to their corresponding β-Tubulin band intensities. The normalized values from the Snail KO cells were plotted relative to those from the corresponding WT control under each condition. (G) Schematic representation of the SARS-CoV-2 entry assay. WT and Snail KO cells were exposed to SARS-CoV-2 for the indicated durations, followed by PBS washing and proteinase K treatment to remove surface-bound virions before sample collection. (H) RT-qPCR quantification of intracellular SARS-CoV-2 *E* RNA in WT and Snail KO cells during the entry assay at the indicated time points. The normalized values from the Snail KO cells were plotted relative to those from the corresponding WT control at each time point. Data are represented as mean ± SEM from three independent biological replicates (n = 3). Each dot represents one biological replicate. Statistical significance was determined using an unpaired Student’s *t*-test. *, **, and *** denote *P* < 0.05, *P* < 0.01, and *P* < 0.001, respectively; n.s., not significant.

Having identified *ACE2* as an upregulated candidate from the transcriptomic analysis, we first validated its expression independently by RT-qPCR. *ACE2* transcript abundance was markedly increased in Snail KO cells as against the WT cells under both infected and mock conditions (Figure 4C), confirming the transcriptional phenotype identified in the RNA seq analysis. We next asked whether this increased *ACE2* mRNA was reflected at the protein level. ACE2 protein abundance was markedly increased in Snail KO cells under basal condition (Figure 4D). A similar difference between WT and Snail KO cells was also observed following SARS-CoV-2 infection, with ACE2 remaining consistently higher in Snail KO cells than in WT cells (Figures 4 E&F). We next asked if this elevated abundance of ACE2 receptor in Snail KO cells translated to higher SARS-CoV-2 entry. Snail KO and WT cells were incubated with SARS-CoV-2 for up to 3 hrs and the cells were collected at every 30 minutes for viral entry assay (Figure 4G). Viral RNA in the collected samples was measured by RT-qPCR. Consistent with their higher ACE2 abundance, Snail KO cells showed higher levels of intracellular SARS-CoV-2 *envelope* (*E*) RNA than WT cells throughout the entire time course (Figure 4H). The difference was apparent from 30 minutes of incubation and remained evident at subsequent time points, supporting enhanced SARS-CoV-2 entry in the absence of Snail. Together, these findings suggest that Snail shapes the broader epithelial entry landscape, in line with the existing understanding in its role in EMT (9). In the context of SARS-CoV-2, increased ACE2 abundance and enhanced viral entry in Snail KO cells link this remodelling to greater viral permissiveness and provide a virus specific example of how Snail dependent epithelial programs may influence viral tropism.

### Snail negatively regulates ACE2 transcription

The inverse relationship between Snail and ACE2 expression suggested that Snail may negatively regulate ACE2. We therefore examined whether restoring Snail expression could reverse the elevated ACE2 phenotype observed following Snail depletion. HA-Snail was expressed in WT and Snail KO cells, and ACE2 abundance was examined. Snail expression significantly reduced ACE2 protein abundance in both WT and Snail KO cells compared with their corresponding empty-vector controls (Figures 5A-C). We next independently examined their relationship during SARS-CoV-2 infection in both the WT and Snail KO cells and observed a similar reduction in ACE2 protein levels following Snail expression in both cellular backgrounds (Figures 5D-F). Consistent with the protein level changes, *ACE2* transcript abundance was also significantly reduced following Snail expression both in the uninfected and SARS-CoV-2 infected WT and Snail KO Huh7 cells (Figure 5 G&H). Together with the increased ACE2 expression observed following Snail depletion, these results confirmed that Snail negatively regulates ACE2.

**Figure 5.**
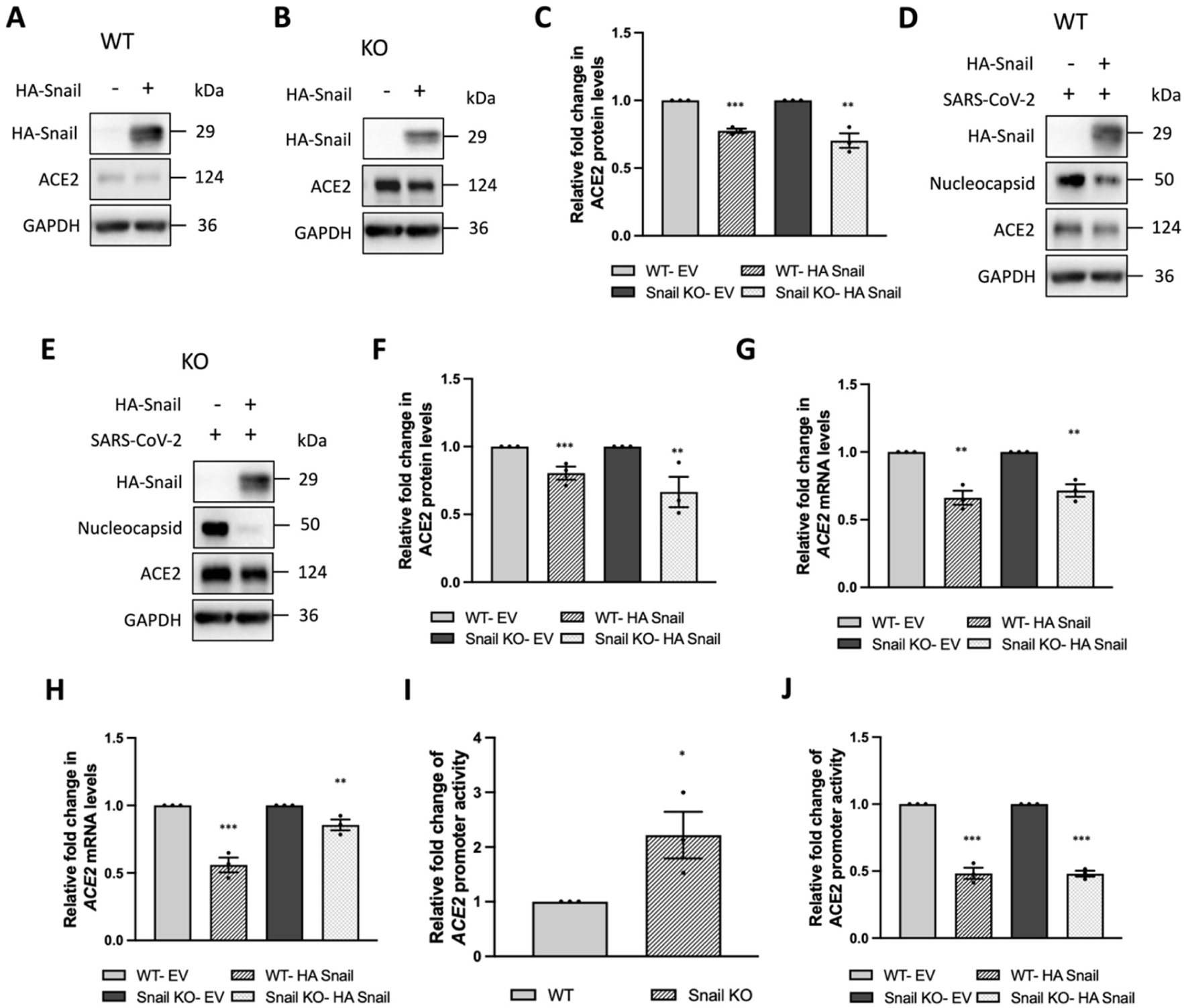
Snail negatively regulates ACE2 transcription. (A) Immunoblot showing HA-Snail, ACE2, and GAPDH protein levels in WT Huh7 cells transfected with pcDNA4/TO empty vector or HA-Snail-pcDNA4/TO under mock conditions. (B) Immunoblot showing HA-Snail, ACE2, and GAPDH protein levels in Snail KO Huh7 cells transfected with pcDNA4/TO empty vector or HA-Snail-pcDNA4/TO under mock conditions. (C) Densitometric quantification of ACE2 protein levels from panels (A) and (B). Individual ACE2 band intensities from pcDNA4/TO empty vector- and HA-Snail-pcDNA4/TO-transfected samples were normalized to their corresponding GAPDH band intensities. The normalized values from the HA-Snail-pcDNA4/TO-transfected samples were plotted relative to those from the corresponding pcDNA4/TO empty vector-transfected control for each cell background. (D) Immunoblot showing HA-Snail, SARS-CoV-2 Nucleocapsid, ACE2, and GAPDH protein levels in SARS-CoV-2-infected WT Huh7 cells transfected with pcDNA4/TO empty vector or HA-Snail-pcDNA4/TO. (E) Immunoblot showing HA-Snail, SARS-CoV-2 Nucleocapsid, ACE2, and GAPDH protein levels in SARS-CoV-2-infected Snail KO Huh7 cells transfected with pcDNA4/TO empty vector or HA-Snail-pcDNA4/TO. (F) Densitometric quantification of ACE2 protein levels from panels (D) and (E). Individual ACE2 band intensities from pcDNA4/TO empty vector- and HA-Snail-pcDNA4/TO-transfected samples were normalized to their corresponding GAPDH band intensities. The normalized values from the HA-Snail-pcDNA4/TO-transfected samples were plotted relative to those from the corresponding pcDNA4/TO empty vector-transfected control for each cell background. (G) RT-qPCR analysis of *ACE2* mRNA expression in WT and Snail KO cells transfected with pcDNA4/TO empty vector or HA-Snail-pcDNA4/TO under mock conditions. Individual *ACE2* transcript levels were normalized to *GAPDH*. The normalized values from the HA-Snail-pcDNA4/TO-transfected samples were plotted relative to those from the corresponding pcDNA4/TO empty vector-transfected control for each cell background. (H) RT-qPCR analysis of *ACE2* mRNA expression in SARS-CoV-2-infected WT and Snail KO cells transfected with pcDNA4/TO empty vector or HA-Snail-pcDNA4/TO. Individual *ACE2* transcript levels were normalized to *GAPDH*. The normalized values from the HA-Snail-pcDNA4/TO-transfected samples were plotted relative to those from the corresponding pcDNA4/TO empty vector-transfected control for each cell background. (I) Luciferase reporter assay comparing basal activity of the distal *ACE2* promoter in WT and Snail KO cells. Firefly luciferase activity was normalized to Renilla luciferase activity, and the normalized promoter activity in Snail KO cells was plotted relative to that in WT cells. (J) Luciferase reporter assay measuring distal *ACE2* promoter activity in WT and Snail KO cells transfected with pcDNA4/TO empty vector or HA-Snail-pcDNA4/TO. Firefly luciferase activity was normalized to Renilla luciferase activity, and the normalized promoter activity in HA-Snail-pcDNA4/TO-transfected cells was plotted relative to that in the corresponding pcDNA4/TO empty vector-transfected control for each cell background. Data are represented as mean ± SEM from three independent biological replicates (n = 3). Each dot represents one biological replicate. Statistical significance was determined using an unpaired Student’s *t*-test. *, **, and *** denote *P* < 0.05, *P* < 0.01, and *P* < 0.001, respectively; n.s., not significant.

Since Snail is an E-box binding transcriptional repressor, we next asked whether it could transcriptionally regulate *ACE2* expression. To address this, we examined two *ACE2* regulatory regions obtained from independent promoter annotations. First, we used the NCBI-derived distal *ACE2* promoter region on chromosome X (NC_000023.11:15600911-15603000), which spans approximately 2.1 kb of the upstream regulatory region. NCBI annotates this interval as an *ACE2* promoter region (38). Sequence analysis of this region identified seven candidate E-box motifs, including two sequences corresponding (E1 & E4) to preferential Snail binding motifs, together with several additional predicted transcription factor binding sites (39) (Supplementary Figure S5A). The NCBI-annotated sequence corresponded to −2089/-1 relative to the ATG start site; however, the distal boundary contained only part of one candidate E-box. We therefore extended the sequence upstream by three nucleotides to −2092/−1 to include the complete motif. This distal *ACE2* promoter fragment was PCR amplified and cloned upstream of the firefly luciferase reporter in the pGL4.1 vector for promoter activity analysis.

We additionally examined a proximal *ACE2* promoter region derived from the Eukaryotic Promoter Database (EPD). Based on the EPD annotation, we generated a proximal *ACE2* promoter construct spanning −454/-1 relative to the ATG start site (40) and cloned this fragment into the same pGL4.1 reporter backbone. In contrast to the NCBI derived region, no canonical E-box motifs were identified within this proximal fragment, although predicted binding sites for GATA6 and HNF1α/β were present (39) (Supplementary Figure S5B). We therefore compared the transcriptional activity of these promoter regions.

We first examined the basal activity of the two promoter constructs. The distal *ACE2* promoter showed clear reporter activity, whereas the proximal *ACE2* promoter showed little measurable activity above the pGL4.1 vector background (Supplementary Figures S 5C&D). We therefore focused subsequent Snail response experiments on the transcriptionally active distal promoter construct. This *ACE2* promoter showed significantly higher basal promoter activity in Snail KO cells than in

WT cells (Figure 5I), suggesting the absence of Snail-mediated repression of *ACE2* promoter in them. Agreeably, overexpression of Snail caused significant and comparable reduction of promoter activity in both WT and Snail KO cells as against their corresponding empty vector controls (Figure 5J). Together, these results suggest that Snail negatively regulates ACE2 at the transcriptional level.

Based on these findings, we propose a model in which Snail limits SARS-CoV-2 entry by maintaining lower ACE2 expression and a relatively restrictive epithelial state (Figure 6). During infection, SARS-CoV-2 progressively reduces Snail protein despite maintained or increased *SNAI1* transcript levels. Loss of Snail increases ACE2 abundance and shifts the cells towards a more permissive epithelial state, thereby modulating SARS-CoV-2 tropism.

**Figure 6.**
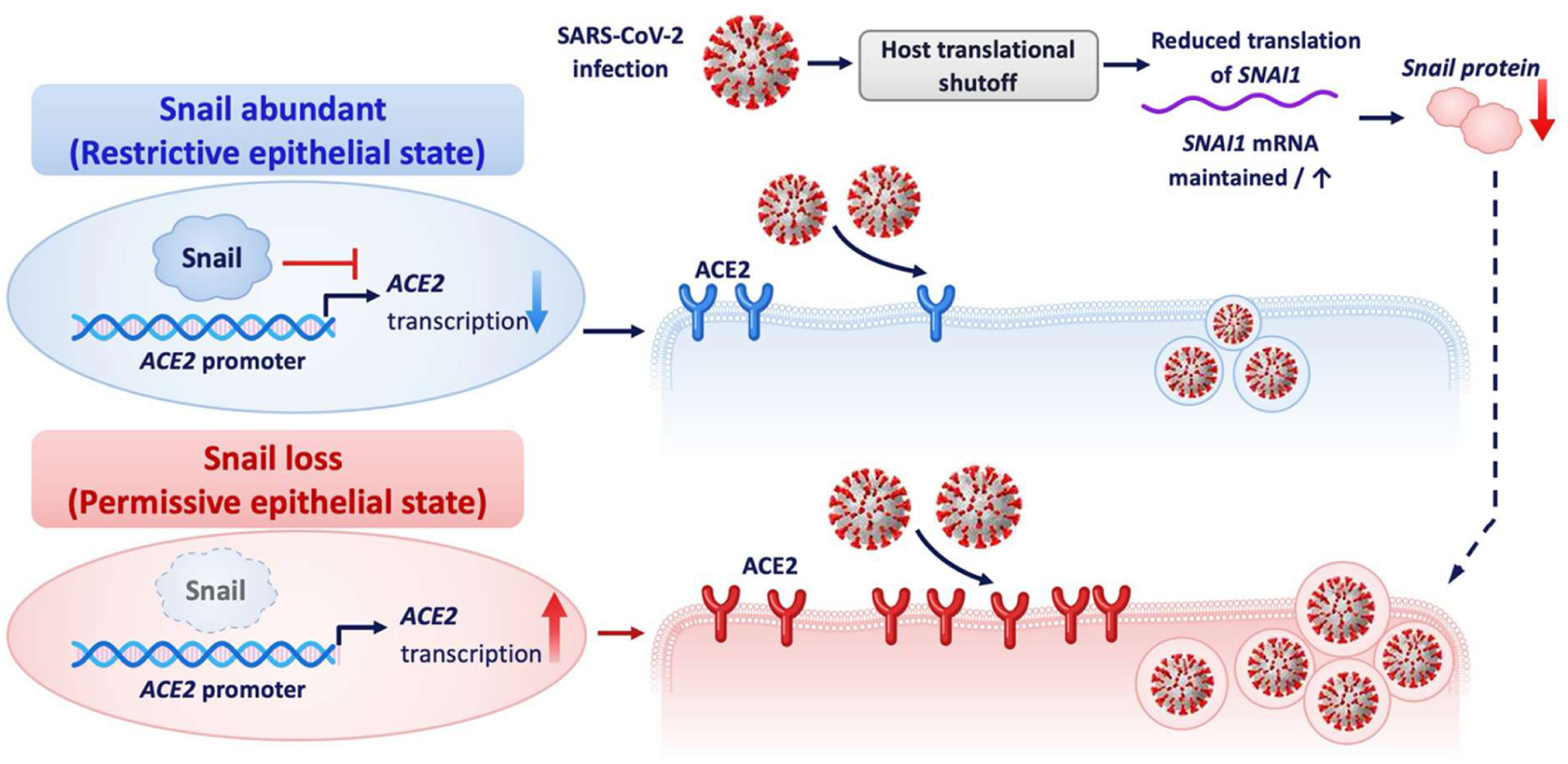
Proposed model of Snail mediated regulation of SARS-CoV-2 permissiveness. Schematic model illustrating the role of Snail in regulating the epithelial entry state during SARS-CoV-2 infection. In Snail expressing cells, Snail negatively regulates *ACE2* transcription, resulting in lower ACE2 abundance and a relatively restrictive epithelial state. Increased *ACE2* transcription following loss of Snail promotes ACE2 surface abundance, establishing a more permissive epithelial state associated with enhanced SARS-CoV-2 entry and infection. Utilizing this regulatory circuit, SARS-CoV-2 infection promotes a more permissive Snail-deficient and ACE2-high condition. SARS-CoV-2-induced host translational shutoff reduces *SNAI1* translation despite maintained or increased *SNAI1* mRNA abundance, leading to depletion of Snail protein. The dashed arrow indicates the proposed contribution of infection associated Snail loss to weakening of the pre-existing restrictive epithelial state during ongoing infection.

## DISCUSSION

Viral tropism reflects the permissiveness of the host system to viral infections and is determined largely by entry receptors and other host-specific factors (6). Cell-specific characteristics and cellular states are regulated by complex interplays between various growth factors and transcription factors (41, 42). It is reasonably well established that stem cells show elevated levels of viral restriction mediated by intrinsic factors (43), underlining the importance of these factors in determining viral permissiveness. Our study places the EMT transcription factor Snail at the interface between epithelial cell state and antiviral restriction and advances two related concepts. First, Snail constitutes a cell-intrinsic restriction factor for SARS-CoV-2 that extends beyond its previously described role in innate immune signaling. We previously showed that RLR activation induces Snail and that Snail restricts RNA-virus infection through antiviral signaling (17). Here, loss of Snail establishes a more permissive epithelial state characterized by altered surface-associated programs, increased ACE2 abundance, and enhanced SARS-CoV-2 entry. This suggests that Snail can shape the epithelial entry landscape and thereby contribute to cellular determinants of viral tropism. Second, SARS-CoV-2 infection uncouples *SNAI1* transcription from Snail protein abundance, allowing an antiviral program to remain transcriptionally active while becoming functionally depleted.

Snail appears to restrict different RNA viruses through distinct mechanisms. Unlike our previous RLR-linked phenotype (17), the restriction observed here was not accompanied by major changes in the type I or III IFN readouts examined, and Snail KO cells retained responsiveness to exogenous IFN-α. Snail also restricted the Delta variant, which has been reported to show greater resistance to IFN-mediated restriction than earlier SARS-CoV-2 isolates (22). Together, these findings suggest that Snail-mediated restriction of SARS-CoV-2 is not primarily dependent on amplification of canonical IFN signaling. Epithelial plasticity can alter viral susceptibility through changes in entry receptors and other surface-associated factors (12, 13). Accordingly, the Snail KO transcriptome showed a substantial, infection-independent program enriched for apical membrane, brush border, and junction-associated genes, consistent with the established role of Snail as an E-box-binding regulator of epithelial identity (15). Such remodelling provides a route by which epithelial cell state could influence the repertoire and organization of host factors encountered by different viruses and thereby contribute to cellular tropism.

Within this broader surface-associated program, *ACE2* was upregulated following Snail loss, linking epithelial remodelling to a key determinant of SARS-CoV-2 entry.

This relationship is consistent with previous studies linking ACE2 expression with epithelial-mesenchymal plasticity, particularly the inverse relationship between ZEB1 and ACE2 (14). Interestingly, negative associations between *SNAI1* and *ACE2* expression have also been reported across several aerodigestive and respiratory datasets (14). Snail could therefore regulate ACE2 through direct or indirect mechanisms. Snail is a well-established E-box binding transcriptional repressor and directly represses epithelial genes such as *CDH1* (15). Snail can also promote ZEB1 expression during EMT (44), providing a potential indirect route for ACE2 regulation, while ZEB1 has been shown to occupy E-box containing regions of the *ACE2* promoter and repress its expression (14). Our findings extend this framework by showing reciprocal regulation of endogenous ACE2 expression and *ACE2* promoter activity following Snail loss and restoration. The NCBI derived distal *ACE2* promoter region that was responsive to Snail contained multiple candidate E-box motifs, including sequences corresponding to preferential Snail-binding motifs. In contrast, the EPD defined proximal *ACE2* promoter lacked canonical E-box motifs and showed little measurable promoter activity under our experimental conditions. The presence of multiple candidate E-boxes within the responsive promoter region provides a possible basis for this regulation, although direct Snail occupancy and the contribution of individual E-box motifs remain to be established.

ACE2 regulation provides a functional connection between the broader epithelial entry landscape controlled by Snail and the SARS-CoV-2 specific phenotype observed here. Increased expression of ACE2, the principal cellular receptor for SARS-CoV-2 (1), in Snail KO cells was accompanied by enhanced viral entry, consistent with Snail limiting cellular permissiveness by restraining receptor availability, although ACE2 is unlikely to be the sole contributor. Thus, the ACE2 entry phenotype provides a virus specific example of how Snail dependent epithelial programs may influence viral tropism. The timing of Snail loss is important in interpreting this relationship. Basal levels of Snail at the time of infection can therefore shape the cellular environment encountered by the incoming virus. However, Snail depletion well after the infection is established, is therefore unlikely to affect entry of the initiating inoculum. Progressive loss of Snail may instead weaken this pre-existing restriction and favor subsequent rounds of viral spread, although this possibility remains to be directly tested. Given that Snail is regulated upstream by many growth factors, it is likely that any of these factors triggered by the initial sites of infection could prepare for systemic resistance to viral infection. The Snail-ACE2 relationship is also biologically double edged. Sustained Snail activity and EMT are associated with loss of epithelial integrity, fibrosis, and malignant progression (9), whereas ACE2, despite functioning as the SARS-CoV-2 receptor, has important physiological and tissue-protective functions (45). Thus, these findings do not support indiscriminate activation of Snail or inhibition of ACE2 as antiviral strategies. Instead, defining specific Snail regulated pathways that maintain a restrictive epithelial state may provide more selective targets.

Snail protein progressively declined during SARS-CoV-2 infection, whereas *SNAI1* transcript levels were maintained and, in some cell types, increased. Previous reports have described *SNAI1* induction and EMT associated changes following SARS-CoV-2 infection or expression of individual viral proteins (16). Our findings show that this transcriptional response is uncoupled from Snail protein abundance during infection. Collectively, our results suggest that the progressive loss of Snail can only be explained by reduced protein synthesis. This is relevant because Snail is intrinsically short-lived and undergoes rapid proteasomal turnover (46). SARS-CoV-2 is well known to suppress host protein synthesis, including through Nsp1 mediated obstruction of the ribosomal mRNA entry channel and additional mechanisms that impair host mRNA expression and translation (32, 47). However, this host shutoff is not absolute, and accumulating evidence indicates that individual cellular transcripts differ considerably in their sensitivity to Nsp1-mediated repression. Ribosome profiling of Nsp1-expressing cells identified a subset of host transcripts that retained relatively high translation efficiency despite the overall reduction in protein synthesis, including transcripts encoding components of the translation machinery, RNA-binding proteins, and proteins involved in host immune responses and other cellular processes (48). Many of these transcripts contained 5′TOP motifs, suggesting that transcript-specific regulatory features can permit preferential translation during Nsp1-mediated shutoff. Consistent with this selectivity, Nsp1 preferentially destabilizes multi-exon host transcripts, whereas intronless transcripts, including interferon genes, show greater resistance to Nsp1-induced RNA degradation (49). This selective resistance may help explain why components of the antiviral response can remain induced at later stages of infection despite broad suppression of host translation. Thus, SARS-CoV-2-induced host shutoff is better viewed as a selective remodelling of the translational landscape rather than a uniform cessation of host protein synthesis. In this context, short-lived regulators such as Snail may be particularly vulnerable when their translational engagement is reduced. Given the antiviral role of Snail, its reduction during infection may also contribute to a more permissive cellular state as infection progresses. More broadly, protein stability may therefore influence how strongly individual host restriction factors are affected during virus induced host shutoff.

While our results have been insightful, several limitations need to be discussed. Our polysome associated fractionation data support reduced translational engagement of *SNAI1* at later stages of infection, although direct measurement of nascent Snail synthesis would further strengthen this conclusion. Likewise, the reciprocal effects of Snail loss and restoration on *ACE2* promoter activity support transcriptional regulation of *ACE2*, while the precise contribution of individual E-box motifs remains to be defined. Functional loss-of-function analyses were performed predominantly in Huh7 cells, and extension of the Snail-ACE2 relationship to additional physiologically relevant epithelial models will be important. Nevertheless, our findings identify a broader role for Snail in antiviral defense than previously recognized. By shaping epithelial surface programs and ACE2 associated entry, Snail contributes to the cellular state that determines SARS-CoV-2 permissiveness and may influence cellular determinants of viral tropism. SARS-CoV-2 infection, in turn, progressively depletes this short-lived restriction factor in association with host translational shutoff, potentially limiting the ability of Snail to sustain this restrictive epithelial state as infection progresses. Thus, a host program that initially limits viral entry is itself weakened during infection. Together, these observations connect epithelial plasticity, viral entry, and host translational control within a common framework for understanding how cell state influences viral infection.

## MATERIALS AND METHODS

### Cells and Virus infections

Caco-2, A549, and Vero cells were obtained from ATCC; Huh7 cells were generously gifted by Ralf Bartenschlager (50). Cells were cultured in Dulbecco’s modified Eagle medium (DMEM) supplemented with 10% FBS (Thermo Fisher) (20% FBS for Caco-2), 1× penicillin-streptomycin cocktail (Thermo Fisher), and 1× non-essential amino acids (Thermo Fisher) and maintained at 37°C in a humidified atmosphere containing 5% CO_2_. Cells were routinely passaged at 70-80% confluency, and mycoplasma contamination was periodically monitored. SARS-CoV-2 (isolate hCoV-19/India/TG-CCMB-L1021/2020, GISAID accession EPI_ISL_458046, PANGO lineage B.1.1.8) and the Delta variant (isolate hCoV-19/India/TG-CCMB-CIA4413/2021, GISAID accession EPI_ISL_2775201, PANGO lineage B.1.617.2) were propagated in Vero cells (CCL-81) and titered by plaque formation assay, as previously described (22). For infection experiments, cells were inoculated with virus at a multiplicity of infection (MOI) of 1 for 3 hrs at 37°C in serum-free medium, after which the inoculum was replaced with complete medium and cells were further incubated until the time of harvesting as required for each experiment; mock-infected cells received an equivalent volume of virus-free medium in place of the viral inoculum.

### Plasmids, transfection, and electroporation

The following plasmids were used in this study: PX459 (Addgene, Cat #62988), which encodes wild-type Cas9 and was used for CRISPR-Cas9 mediated generation of Snail KO Huh7 cells; pcDNA4/TO (Thermo Fisher), used as the backbone for N-terminally HA-tagged human Snail (HA-Snail) under a CMV promoter and, in its empty form, as the vector control in all overexpression experiments (17); and pGL4.1 (Promega), used to generate the ACE2 promoter-luciferase reporter construct.

The PX459-sgRNA construct was delivered by transfection with Lipofectamine 3000 (Thermo Fisher, Cat #L3000015) in Opti-MEM according to the manufacturer’s protocol. For HA-Snail overexpression and ACE2 promoter-luciferase reporter co-transfection experiments, plasmid DNA was delivered by electroporation using a Bio-Rad Gene Pulser (0.2 cm gap cuvette, 100 µL Opti-MEM) with a single square-wave pulse of 85 V for 40 ms, using 3 µg total plasmid DNA per 1 × 10⁶ cells (51).

### Generation of Snail KO Huh7

Snail KO Huh7 cells were generated using the CRISPR-Cas9 system. A single-guide RNA (sgRNA) targeting exon 1 of the *SNAI1* gene (5′-AGAGCGCGGCATAGTGGTCG-3′, targeting the antisense strand immediately upstream of a 5′-AGG-3′ protospacer-adjacent motif) was cloned into the PX459 vector, which encodes wild-type Cas9 (Addgene, Cat #62988), using the annealed oligonucleotide pair 5′-CACCGAGAGCGCGGCATAGTGGTCG-3′ (forward) and 5′-AAACCGACCACTATGCCGCGCTCTC-3′ (reverse). The PX459-sgRNA construct was delivered by transfection with Lipofectamine-3000 (Thermo Fisher) in Opti-MEM according to the manufacturer’s protocol. Following transfection, cells were selected with puromycin; the puromycin-selected population was expanded as a pool, from which single clones were subsequently isolated by clonal selection. Knockout validation was performed by immunoblotting to confirm the loss of Snail protein.

### Treatments

#### MG132 treatment

Huh7 cells were infected with SARS-CoV-2 as described above. At 16, 40 or 64 hrs post-infection (hpi), the medium was replaced with fresh medium containing 10 µM MG132 or an equivalent volume of DMSO as vehicle (Invivogen, Cat #tlrl-MG132-2), and cells were harvested 8 hrs after treatment, corresponding to 24, 48, or 72 hpi, respectively. Proteasome inhibition was confirmed by immunoblotting for the accumulation of ubiquitinated proteins.

#### IFN-α treatment

For IFN-α treatment, cells were serum-starved at 70-75% confluency in serum-free DMEM (SFD) for 3 hrs. SFD was then replaced with fresh SFD containing 1,000 U/mL IFN-α (PBL Assay Science, Cat #11100) for 1 hr. Cells were subsequently infected with SARS-CoV-2 for 3 hrs and further incubated in fresh complete DMEM containing either phosphate-buffered saline (PBS; vehicle) or 1,000 U/mL IFN-α for the subsequent 21 hrs. Cells and culture supernatants were harvested at 24 hpi for downstream analyses.

### Cycloheximide chase assay

Huh7 cells were mock-infected or infected with SARS-CoV-2 at an MOI of 1. Prior to harvesting at 30 hpi, cells were treated with DMSO or 100 µg/mL cycloheximide (CHX) (Sigma-Aldrich, Cat #C1988) for the indicated durations (0, 15, 30, 60, 90, or 120 minutes), with a time-matched, CHX-untreated control included at each time point. Cells were harvested by scraping and processed for immunoblotting.

### Polysome fractionation

Huh7 cells were mock-infected or infected with SARS-CoV-2 and lysed at the indicated time points in polysome lysis buffer (20 mM Tris-HCl, pH 8.0, 140 mM KCl, 1.5 mM MgCl2, 1% Triton X-100, 0.5 mM DTT (Sigma-Aldrich, Cat #D9779), 100 µg/mL cycloheximide, 1 × protease inhibitor cocktail (Roche, Cat #11836170001), 1 mg/mL heparin (HiMedia, Cat #TC138), and 100 U RiboLock RNase inhibitor (Thermo Fisher, Cat #EO0381)). Clarified lysates were layered onto a 30% sucrose cushion prepared in the same buffer and subjected to ultracentrifugation at 100,000 × g for 2 hrs at 4°C to sediment polysome-associated material (pellet), separating it from the non-polysomal (supernatant) fraction, using a cushion approach related to that described previously for pelleting ribosome-bound reporter mRNA (52). RNA was extracted from the pellet and supernatant fractions using TRIzol reagent and reverse-transcribed as described in the upcoming section. Transcript association with the polysome-enriched pellet fraction was calculated as ΔCt = Ct(pellet) − Ct(supernatant) and converted to fold enrichment (2-ΔCt).

### Immunoblotting

Cell pellets were lysed in 1 × Nonidet P-40 lysis buffer (1% Nonidet P-40, 50 mM Tris-HCl, 150 mM NaCl (pH 7.5), 1 mM sodium orthovanadate, 10 mM sodium pyrophosphate, and 1 mM PMSF), incubated on ice for 25 minutes with intermittent vortexing, and centrifuged at 15,000 rpm for 30 minutes at 4°C. The protein-containing supernatants were collected and quantified using BCA reagents (Pierce, Cat #23225). Lysates were mixed with Laemmli buffer and were resolved by SDS-PAGE and transferred for immunoblotting with the following primary antibodies: anti-Snail (Cell Signaling Technology, Cat #3879, 1:1000), anti-HA (Cell Signaling Technology, Cat #3724, 1:1000), anti-SARS-CoV-2 Nucleocapsid (Thermo Fisher, Cat #MA5-36272, 1:3000), anti-ACE2 (Thermo Fisher, Cat #MA5-32307, 1:1000), anti-phospho-STAT1 (Cell Signaling Technology, Cat #14994, 1:1000), anti-total STAT1 (Cell Signaling Technology, Cat #9167, 1:1000), anti-IFIT1 (Cell Signaling Technology, Cat #14769, 1:1000), anti-ubiquitin (Cell Signaling Technology, Cat #43124, 1:1000), anti-GAPDH (Cell Signaling Technology, Cat #5174, 1:5000), and anti-β-tubulin (Cell Signaling Technology, Cat #2146, 1:5000). Horseradish peroxidase (HRP)-conjugated secondary antibodies were from Jackson ImmunoResearch, Goat anti-rabbit HRP conjugated (Cat #115-035-146) and Goat anti-mouse HRP conjugated (Cat #115-035-003). Blots were developed by chemiluminescence on a Bio-Rad Chemidoc MP system using Clarity Western ECL substrate (Biorad, Cat #1705061) and WesternBright^TM^ Sirius (advansta, Cat #K-12043-D20) chemiluminescent substrate kits, and band intensities were quantified by densitometry (ImageJ) and normalized to the corresponding loading control.

### RNA extraction, cDNA synthesis, and RT-qPCR

Total cellular RNA was extracted using the NucleoSpin RNA kit (Macherey-Nagel, Cat #740902). Equal amounts of RNA were reverse-transcribed into cDNA using the PrimeScript 1st Strand cDNA Synthesis Kit (TaKaRa, Cat #6110), following the manufacturer’s protocol. Real time quantitative PCR was performed using TB Green Premix (TaKaRa, Cat #RR820) on a LightCycler 480 instrument (Roche) with gene-specific primers, as previously described (17, 21, 22). Transcript levels were normalized to GAPDH. qRT-PCR to quantify SARS-CoV-2 RNA was performed on a Roche LightCycler 480 using an nCOV-19 reverse transcription-PCR (RT-PCR) detection kit (Q-Line Molecular, Cat #COVIDM96PS). The relative fold changes were calculated using the 2^-ΔΔCT method. The following primers were used for qRT-PCR:

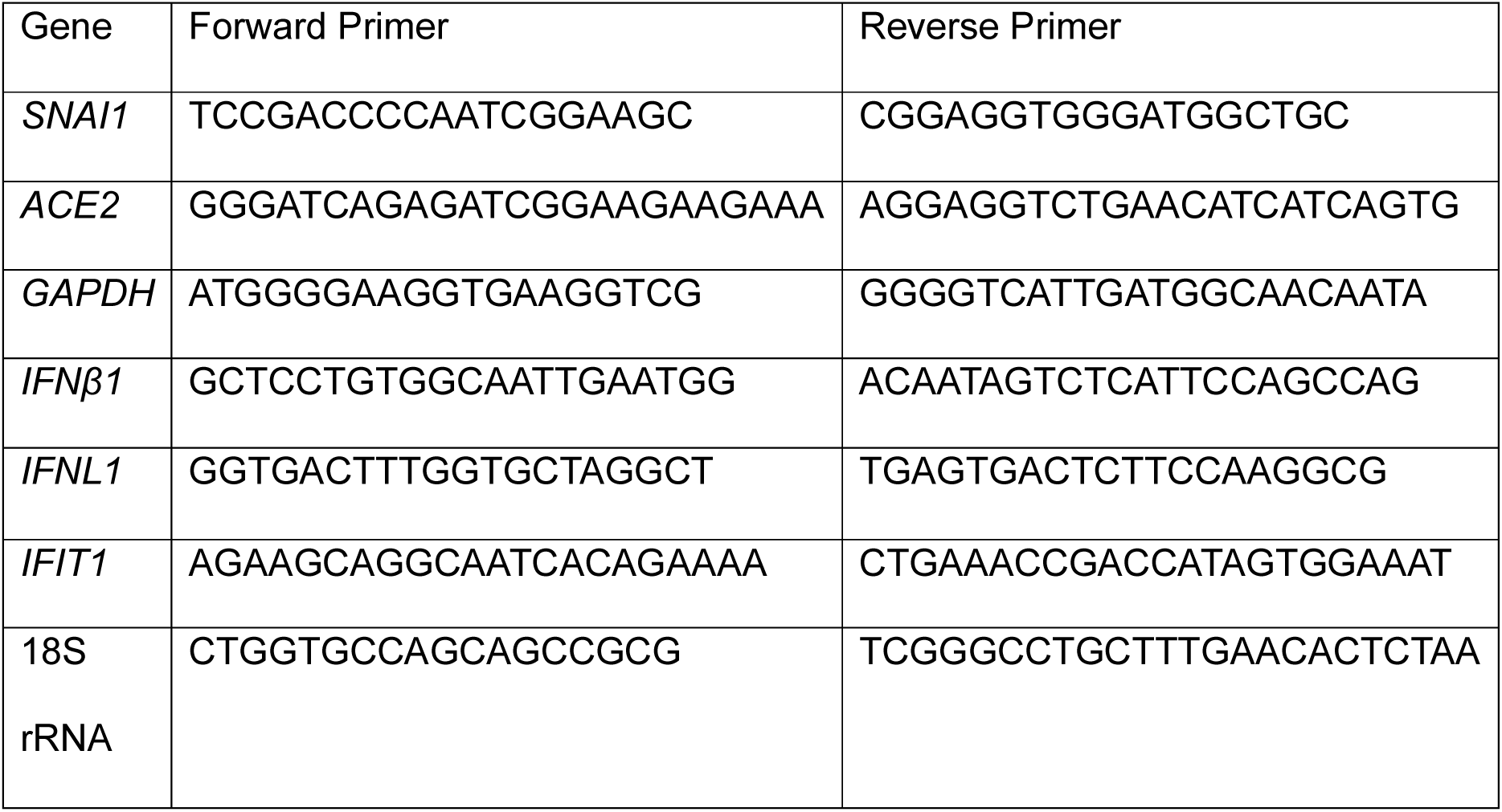

### Northern blotting

Northern blotting for *SNAI1* and *ACTB* transcripts was performed using the DIG Northern Starter Kit (Merck/Roche Cat #12039672910) according to the manufacturer’s instructions. A digoxigenin (DIG)-labeled antisense RNA probe against *SNAI1* was generated by in vitro transcription with T7 RNA polymerase from a PCR template amplified using a forward primer (5′-AAGATGCACATCCGAAGCCAC-3′) and a T7 promoter-tailed reverse primer (5′-TAATACGACTCACTATAGGGGATCATCAAAGTCCTGTGGGGC-3′), yielding a probe corresponding to a fragment of the *SNAI1* coding sequence. Total RNA was resolved on a denaturing gel, transferred to a nylon membrane, and hybridized with the DIG-labeled probe; hybridized probe was detected per the DIG Northern Starter Kit detection protocol.

### RNA deep-sequencing and bioinformatics analysis

RNA-seq library preparation was performed using the MGIEasy RNA Library Prep Set (MGI) according to the manufacturer’s instructions. Briefly, 500 ng of total RNA was used as starting material, and rRNA was depleted using the Ribo-Zero Plus rRNA Depletion Kit (Illumina). The rRNA-depleted samples were fragmented and reverse-transcribed, followed by second-strand synthesis. The resulting DNA was purified using the DNA clean beads provided with the kit, followed by end repair and A-tailing. Barcoding and adaptor ligation were subsequently performed, followed by purification and amplification using adaptor-specific primers. Libraries were quantified using the Qubit dsDNA High Sensitivity Assay Kit (Thermo Fisher Scientific), and fragment size distribution was assessed using a 4200 TapeStation (Agilent Technologies). The libraries were denatured and converted into single-stranded circular DNA, followed by rolling-circle amplification to generate DNA nanoballs. The resulting libraries were loaded onto flow cells, and paired-end sequencing was performed on the MGI 2000 platform (MGI) with a read length of 100 bases.

Data processing and functional enrichment analysis were performed as previously described (21). Briefly, reads were trimmed with Cutadapt to remove MGI adapters and low-quality sequences (Q < 20 or length < 36 bp), then aligned to the human genome (GRCh38) with HISAT2, and the uniquely mapped reads were counted with featureCounts using Ensembl annotation. Genes with fewer than 10 total counts across all samples were removed, and differential expression was assessed with DESeq2 (adjusted *P* < 0.05, |log2FC| ≥ 0.58). GO enrichment was performed with clusterProfiler, and redundant terms were merged using the simplify function (similarity cut-off 0.5, Wang method), after which the top enriched terms were visualized.

### SARS-CoV-2 entry assay

Cells were infected with SARS-CoV-2 at an MOI of 2 and harvested at the indicated time points. Prior to harvesting, cells were treated with proteinase K to digest virions that were surface-bound but had not been internalized, leaving only intracellular (internalized) virus intact. Total RNA was then extracted from the treated cells, reverse-transcribed into cDNA, and intracellular SARS-CoV-2 *envelope* (E) gene RNA was quantified by RT-qPCR (Q-Line Molecular, Cat #COVIDM96PS).

### Luciferase reporter assay

The NCBI-derived distal human ACE2 promoter region (NC_000023.11:15600911–15603000) corresponds to −2089/-1 relative to the ATG start codon. The cloned fragment was extended by three nucleotides at the upstream boundary to −2092/-1 to include the complete boundary E-box motif and was inserted upstream of the firefly luciferase reporter in the pGL4.1 vector. This construct is referred to here as the distal *ACE2* promoter. An additional EPD-defined proximal *ACE2* promoter fragment spanning −454/-1 relative to the ATG start codon was cloned into the same reporter backbone. Cells were co-transfected with the ACE2 promoter-luciferase reporter construct together with pcDNA4/TO empty vector or HA-Snail-pcDNA4/TO, and luciferase activity was measured using the Dual-Luciferase Reporter Assay System (Promega, Cat #E1910) according to the manufacturer’s instructions. Firefly luciferase activity was normalized to Renilla luciferase activity as an internal transfection control.

### Statistics

Statistical significance was assessed by an unpaired Student’s t-test using GraphPad Prism software. The resultant p values were represented as *, **, *** indicating p values ≤ 0.05, 0.01, and 0.001, respectively. All experiments were performed with three biological replicates (n = 3) unless otherwise indicated, and data are presented as mean ± SEM.

### Biosafety

Institutional biosafety clearance was obtained for all experiments involving SARS-CoV-2, which were conducted in the biosafety level 3 (BSL-3) facility at the CSIR-Centre for Cellular and Molecular Biology (22).

## Data availability

All data pertaining to this manuscript are included within the manuscript. The RNA sequencing (RNA-seq) data have been submitted to the Gene Expression Omnibus (GEO) database and are available under the accession number GSE342875.

## ACKNOWLEDGMENTS

We thank Dr. Pavithra L. Chavali for sharing the PX459-Snail Exon-1 plasmid. Special appreciation goes to Mohan Singh Moodu and Amit Kumar for their assistance with experimental logistics. We thank Pragada Rajesh, and Celasta Joby for their assistance with KO cell generation and immunoblotting experiments.

This work was supported by funding from the Department of Biotechnology (DBT), Government of India (BT/PR31038/BRB/10/1798/2019); and National Laboratories Scheme under the ULIP sub-scheme [MLP002604]) to K.H.H. P.P.S received fellowships from the Council of Scientific and Industrial Research (CSIR), Government of India. V.S. received a fellowship from the DBT, Government of India.

## AUTHOR CONTRIBUTIONS

P.P.S. and K.H.H. conceived and designed this study and experiments. P.P.S. performed the experiments. V.S. analyzed the transcriptomics data. P.P.S. and K.H.H. wrote the manuscript with inputs from V.S.

During the preparation of this work, the authors used ChatGPT to improve language and readability. After using this tool/service, the authors reviewed and edited the content as needed and take full responsibility for the content of the publication.

